# A population genomic resource for *Drosophila pseudoobscura*

**DOI:** 10.64898/2026.02.02.703370

**Authors:** Yesbol Manat, Zhao Zheng, Camryn A. Kritzell, Christopher A. Gonzales, Richard P. Meisel

**Affiliations:** Department of Biology and Biochemistry, University of Houston, Houston, TX 77204; Current affiliation: Department of Biostatistics and Data Science, University of Texas Health Science Center, Houston, TX 77030; Current affiliation: Center for Depression Research and Clinical Care, University of Texas Southwestern, Dallas, TX 75390

**Keywords:** chromosomal inversions, population structure, Tajima’s D, neo-sex chromosomes

## Abstract

*Drosophila pseudoobscura* is an historically important organism in evolutionary genetics, serving as a model system in studies of chromosomal inversions, speciation, sex chromosome evolution, and sex-ratio drive. However, previous population genetics analysis of *D. pseudoobscura* focused on individual chromosomes or used fragmented genome assemblies as a reference. To address these shortcomings, we generated a *D. pseudoobscura* population genomics resource consisting of newly sequenced genomes from 60 inbred lines sampled across the species’ geographic range in North America. Using these data and a chromosome-scale reference genome, we examined patterns of nucleotide diversity and population structure across the chromosomes. We found no strong evidence of population structure on most chromosomes, consistent with prior results. In contrast, we identified population structure on the third chromosome, which we attributed to a well-characterized inversion polymorphism. We assigned individual third chromosome haplotypes to inversion arrangements, demonstrating how tests for population structure can be used to identify polymorphic chromosomal rearrangements. Tajima’s *D* was negative across most of the genome, consistent with a recent population expansion. However, the distribution of genetic variation differed across third chromosome inversion arrangements in ways that were consistent with their hypothesized evolutionary histories, and we identified inter-arrangement genetic differentiation that could be attributed to the inversions suppressing genetic exchange. The population genomic data we have collected is publicly available and will support future research on evolutionary genetics.

**Summary:** The genomes of 60 isolates of *Drosophila pseudoobscura* were sequenced and analyzed. This species is a model organism for multiple areas in evolutionary genetics research, including chromosomal rearrangement, sex chromosomes, and speciation. This article presents the largest population genomic data set collected in this species. The analysis of the data demonstrates how population structure detection approaches can be used to identify polymorphic chromosomal inversions. The data presented will be valuable for future work on fundamental questions in population genetics.

## Introduction

Population genetics processes are greatly influenced by the structural organization of the genome. For example, chromosomal inversions suppress recombination in heterozygotes, allowing for the maintenance of linkage disequilibrium amongst coadapted complexes of alleles (i.e., supergenes) that would otherwise be broken apart by recombination (Wellenreuther and Bernatchez 2018; Berdan et al. 2023). Similarly, Robertsonian fusions between chromosomes can affect the recombination landscape, thereby shaping large-scale patterns of genetic diversity (Guerrero and Kirkpatrick 2014; Cicconardi et al. 2021; Yoshida et al. 2023). Because of these effects, both chromosomal inversions and fusions have been implicated as important contributors to the speciation process (White 1969; Rieseberg 2001; Faria and Navarro 2010). Fusions between sex chromosomes and autosomes may have especially pronounced effects because they cause ancestrally autosomal regions of the genome to become sex-linked, thereby dramatically shifting the effects of sex-specific selection pressures and hybridization between divergent karyotypes (Zhou and Bachtrog 2012; Bracewell et al. 2017; Wang et al. 2022).

*Drosophila pseudoobscura* is a classic model organism for exploring the effects of genomic rearrangements on genetic diversity, adaptation, and speciation. *D. pseudoobscura* harbors a rich inversion polymorphism on its third chromosome, which was one of the first population genetic data sets ever analyzed (Dobzhansky and Sturtevant 1938; Anderson et al. 1975; Anderson et al. 1991). In addition, *D. pseudoobscura* has been widely used to study the role of chromosomal inversions in species divergence and reproductive isolation (Kulathinal et al. 2009; Fuller et al. 2019; Korunes et al. 2021). Furthermore, the Y and X chromosomes both have features that make them uniquely informative about sex chromosome evolution. Many genes that are Y-linked in other *Drosophila* species are now found on the diminutive chromosome 5 (“dot” chromosome, or element F) in *D. pseudoobscura*, and the species has a new Y chromosome not found beyond its closest relatives (Carvalho and Clark 2005; Larracuente et al. 2010; Bracewell and Bachtrog 2020). There was also a fusion between the ancestral *Drosophila* X chromosome and an autosome in the evolutionary lineage leading to *D. pseudoobscura*, which created a neo-X chromosome that is used as a natural experiment for studying the effects of X-linkage on molecular evolution (Sturgill et al. 2007; Vicoso et al. 2008; Meisel et al. 2010; Meisel et al. 2012; Avila et al. 2014). The *D. pseudoobscura* X chromosome also harbors one of the first ever described sex-ratio distortion systems, in which an X chromosome with multiple inversions causes the elimination of Y-bearing sperm (Sturtevant and Dobzhansky 1936; Novitski et al. 1965; Fuller et al. 2020). Moreover, *D. pseudoobscura* has a closely related subspecies (*D. pseudoobscura bogotana*) and a sister species (*D. persimilis*), and this triad is a model system for speciation research, including investigating how chromosomal inversions, X-linked genes, and sex ratio distorters affect the speciation process (Fuller et al. 2019).

To improve the utility of *D. pseudoobscura* as an evolutionary genetics model organism, we sequenced and analyzed the genomes of 60 strains derived from independently isolated wild-caught female flies sampled across most of the species’ range in North America (Figure 1). Previously collected population genomic data sets from *D. pseudoobscura* either only sampled single chromosomes (Fuller et al. 2017; Fuller et al. 2020) or performed genomewide analyses on a total of 31 unique strains (McGaugh et al. 2012; Samuk et al. 2020; Korunes et al. 2021). In addition, prior analyses did not use a chromosome-scale reference genome, which limited their ability to study the distribution of genetic diversity along each chromosome. Our genome-wide data have nearly twice the sample size of all previous genome-wide population genomics analyses, which we analyzed using a chromosome-scale reference. We used our data to identify single nucleotide polymorphisms (SNPs) across all five chromosomes. We then tested for signatures of genetic differentiation between third chromosome inversion arrangements and differences in evolutionary patterns between the X chromosome and autosomes. Our new dataset will allow for additional exploration of genome-wide genetic diversity in *D. pseudoobscura* to address other fundamental questions in evolutionary genetics.

**Figure 1.**
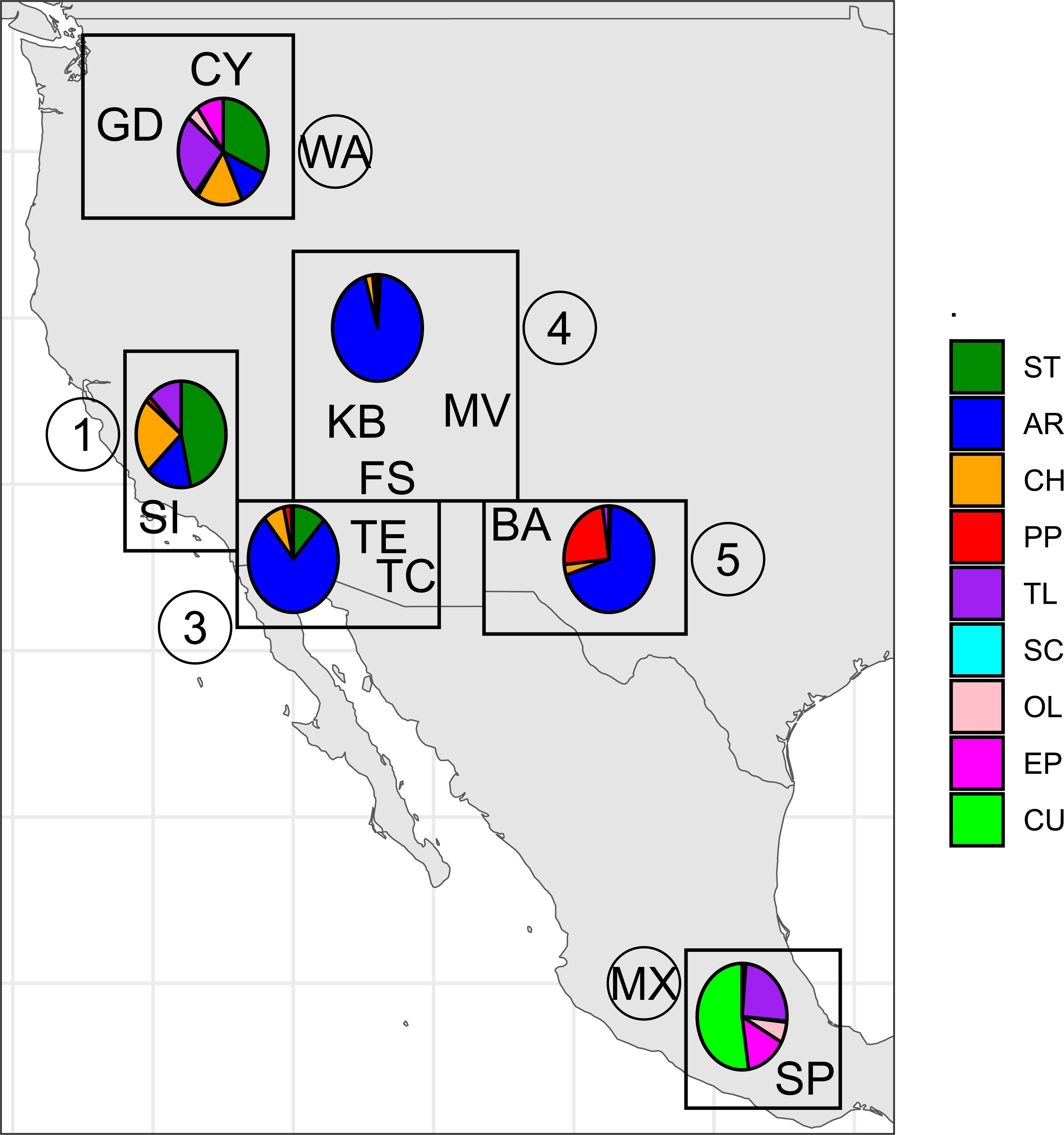
Frequencies of third chromosome arrangements in each sampled population. Each of the 10 sampled populations is shown with its two letter code, with the position of the letters indicating the location of the population (Supplementary Table S1). Each population was assigned to one of six niches, which are represented by rectangles and labeled by numbers or letters within circles (WA, 1, 3, 4, 5, or MX). The frequencies of nine third chromosome arrangements within each niche are represented by pie charts. The evolutionary relationships of the nine arrangements (along with a hypothetical ancestral arrangement, HY) are shown in the top right of the figure. Each arrow represents a single chromosomal inversion event that differentiates adjacent arrangements in the network.

## Materials and Methods

### Drosophila pseudoobscura genome organization

*Drosophila* genomes are organized into five large chromosome arms (Muller elements A–E) and a small “dot” chromosome with fewer than 100 genes (element F) (Muller 1940; Schaeffer et al. 2008). Element A is the ancestral X chromosome of the *Drosophila* genus. Element D was autosomal in the common ancestor of the genus, but it became fused with element A in the lineage leading to *D. pseudoobscura*. We refer to element D as the neo-X chromosome. Element C corresponds to chromosome 3 of *D. pseudoobscura*, which harbors a rich inversion polymorphism consisting of at least 30 different arrangements (Dobzhansky and Sturtevant 1938; Anderson et al. 1991; Wallace et al. 2011). Elements B and E correspond to *D. pseudoobscura* chromosomes 4 and 2, respectively (Schaeffer et al. 2008). *D. pseudoobscura* chromosome 5 consists of element F, which is highly heterochromatic and gene-poor across the *Drosophila* genus (Riddle and Elgin 2018), as well as ancestral Y chromosome genes that transposed to element F after the divergence of the *pseudoobscura* subgroup from sibling lineages (Larracuente et al. 2010; Bracewell and Bachtrog 2020). The *D. pseudoobscura* Y chromosome differs from the Y in other *Drosophila* species, likely because of processes related to the X-autosome fusion and Y-autosome transpositions (Carvalho and Clark 2005; Carvalho et al. 2009). We did not include the Y chromosome in our analysis (because it is not present in the reference genome assembly), but we did include the ancestral X, neo-X, and all autosomes.

### Strain origins and sampling strategy

We sampled 64 strains of *D. pseudoobscura* that were each generated from single females collected from across the species range (Figure 1). These isofemale strains were supplied by the *Drosophila* Species Stock Center. We have listed all of the information the stock center could provide about each strain in Supplementary Table S1, including the strain IDs. Each strain was sampled from one of 10 populations, with each population represented by 1–18 isofemale strains. Each of the 10 populations can be assigned to one of six ecological niches, which differ in climate and third chromosome arrangement frequencies (Dobzhansky 1948; Anderson et al. 1991; Schaeffer 2008). Only 60 samples were retained after filtering for sequencing coverage (see below). VCF files and detailed analysis methods are available in the Texas Data Repository (Manat and Meisel 2026).

### Sequencing methods and sample preparation

We extracted DNA from 5–10 individual flies from each of the 64 strains using a phenol:chloroform protocol (Sambrook and Russell 2006). The DNA was used to construct Illumina sequencing libraries using the Nextera® XT DNA Sample Preparation Kit. All 64 libraries were sequenced on four separate runs of a NextSeq500 instrument at the University of Houston Seq-N-Edit core, with 150 bp paired-end reads.

### Variant calling and genotyping

We used a combination of BWA, GATK, and FreeBayes to determine the genotypes of each of the strains that we sequenced. We first mapped the sequencing reads to *D. pseudoobscura* genome assembly UCI_Dpse_MV25 (GCA_009870125; Liao et al. 2021) using BWA-MEM (Li 2013). Sequencing coverage statistics were calculated from BAM files using *samtools coverage* implemented in SAMtools v1.18 (Danecek et al. 2021), and mean read depth was estimated for each chromosome in each strain (Supplementary Figure S1). We next used GATK v4.0.8.1 to identify duplicate reads, call variants, and recalibrate base calls based on high quality variants that we had identified (McKenna et al. 2010; Poplin et al. 2018; Van der Auwera and O’Connor 2020). We then used FreeBayes v1.3.7 (Garrison and Marth 2012) to genotype all 60 strains separately for each scaffold, providing population assignments with the - *-populations* option to partition the Bayesian inference model according to the populations we sampled (Figure 1). To optimize runtime, the processes were carried out separately on 2 Mb segments of each chromosome (except for chromosome 5, which is only 1.88 Mb in length), and then we merged the VCFs using BCFtools concat (Danecek et al. 2021). Variants were filtered to retain only those with less than 50% missing data (--max-missing 0.5), with a minimum phred quality score of 30 (--minQ 30) and a read depth of 3 (--minDP 3) using VCFtools 1.18-GCC-12.3.0 (Danecek et al. 2011). This left us with 2,105,265 SNPs across the entire genome (Supplementary Table S2). We finally removed strains with more than 50% missing genotypes at variable sites, which excluded 4 strains and reduced the dataset from 64 to 60 strains (Supplementary Table S1).

### Population structure analysis

We used the data from the 60 retained strains to test for population structure using principal component analysis (PCA) and fastSTRUCTURE v1.0 (Raj et al. 2014). Prior to performing these analyses, we used VCFtools 1.18-GCC-12.3.0 (Danecek et al. 2011) to exclude variants with a minor allele frequency <0.05 (--maf 0.05) and a genotype call rate <95% (--max-missing 0.95), thereby removing rare and poorly genotyped variants. These filters substantially reduced the number of retained SNPs per chromosome, with an average retention rate of 39.5% across all chromosomes after filtering (Supplementary Table S2). We used the 831,808 remaining SNPs for population structure analysis. First, we performed PCA with *PLINK v2* (Chang et al. 2015), using linkage disequilibrium-pruned SNPs identified with the*--indep-pairwise 200 25 0.4* option.

We also used fastSTRUCTURE to identify how many population clusters (*K*) best explained the data for each chromosome. Our initial runs of fastSTRUCTURE used a simple (flat beta) prior, and we compared the fit of models 1 *≤ K ≤* 10. Estimating the most appropriate value of *K* from genotype data is challenging (Pritchard et al. 2000). We used the chooseK.py script in fastSTRUCTURE, which provides two separate approaches to identify the best fitting *K* value to the data: one that maximizes marginal likelihood and another that best explains the structure in the data (Raj et al. 2014). If these two estimates of the best fitting *K* values were equal for a chromosome, we used that value of *K* for that chromosome. However, if the two values of *K* differed for a given chromosome, we re-ran fastSTRUCTURE with a logistic prior using *K* values within the range of the two previously identified best fitting values, following the recommended approach (Raj et al. 2014). We used the minimum *K* from the two approaches using the logistic prior as our *K* value for that chromosome. We then used the selected *K* for each chromosome to probabilistically assign each of the 60 strains to populations for each chromosome. We used these population assignments to assign some third chromosome haplotypes to inversion arrangements based on previously collected karyotype information (Fuller et al. 2016).

### Genetic variation across chromosomes

We calculated population genetic statistics on the data prior to filtering for minor allele frequencies and genotype call rates. These analyses were performed on data without the minor allele frequency filter applied because population genetics metrics are sensitive to the frequencies of rare alleles. To gain insights into the evolutionary forces acting on different parts of the genome, we separately analyzed data from intergenic, intronic, and synonymous sites, which may be subject to distinct selective pressures (Ellegren and Galtier 2016). To those ends, we assigned the 2,105,265 SNPs (Supplementary Table S2) to one of four functional categories (intergenic, intronic, synonymous, or missense) using SnpEff v5.2e (Cingolani, Platts, et al. 2012; Cingolani, Patel, et al. 2012).

We also separately analyzed the third chromosome data within subsets of strains that we assigned to specific inversion arrangements. In these arrangement-specific analyses, we used previous mapping information to determine the locations of the chromosome 3 inversion breakpoints and to convert the coordinates of the reference genome (which carries an AR arrangement) into the order found in the TL arrangement (Wright 2023).

The following metrics were calculated within 100 kb windows with 10 kb step sizes. Tajima’s *D* statistic (Watterson 1975; Tajima 1989) was calculated using VCFtools v0.1.16 and VCF-kit (Danecek et al. 2011; Cook and Andersen 2017) for intergenic, intronic, and synonymous sites across the *D. pseudoobscura* genome. Tajima’s *D* measures the difference between between the average pair-wise diversity (*π*) and the number of segregating sites (*Θ*_W_). We therefore also separately calculated *π* and *Θ*_W_ using VCFtools v0.1.16 across the sames sits as Tajima’s *D*. Lastly, Weir and Cockerham’s F_ST_ (1984) was calculated for intergenic sites between third chromosome arrangement haplotypes using the weighted method implemented in VCFtools v0.1.16. All calculations were performed within the same window and step sizes to ensure comparability across statistics.

## Results

We analyzed genome sequence data collected from 60 *D. pseudoobscura* strains that were sampled from 10 populations across the species range in North America (Figure 1) (Supplementary Table S1). Mean sequencing depth per strain was 61.5, ranging from 5.8 to 425 across chromosomes and strains (Supplementary Figure S1). We used these sequencing data to call variants and genotype each of the strains.

### Chromosome-specific population structure

We tested for population stratification in our data using fastSTRUCTURE (Raj et al. 2014). To those ends, we identified how many population groups (*K*) could best explain the genetic diversity on each chromosome. We first used a simple (flat beta) prior and *K* values ranging from 1 to 10, assessing model fit using two criteria (see Methods). The two criteria for the best fitting *K* value were consistent for chromosome 2 (*K*=2) and chromosome 3 (*K*=5) (Supplementary Table S3). However, for chromosome 2, nearly all (56/60) of the strains had >99.999% assignment to one of the populations, and an additional strain had >95% assignment to that same population (Figure 2A). Only three strains had >5% assignment to a second population, and only two of those had >35% assignment to the second population. Three of the four strains with the highest proportional assignment to the second population were derived from females sampled from Mexico (SP population), but most of the Mexico-derived strains had >99% assignment to the first population. The fourth strain in the second population was from Niche 5 (BA population). The SP and BA populations were on the southern end of our sampling area (Figure 1). Consistent with the fastSTRUCTURE results, PCA of chromosome 2 showed little evidence of population structure (Supplementary Figure S2). The first principal component (PC1) explained 13.47% variation, with one sample each from the BA and SP populations separated from the main cluster along the PC1 axis. Also consistent with fastSTRUCTURE, the majority of samples from the BA and SP populations clustered with the other strains. We thus conclude that, while there was some evidence for population structure along the north-south axis of the species range for chromosome 2, the evidence was weak.

**Figure 2.**
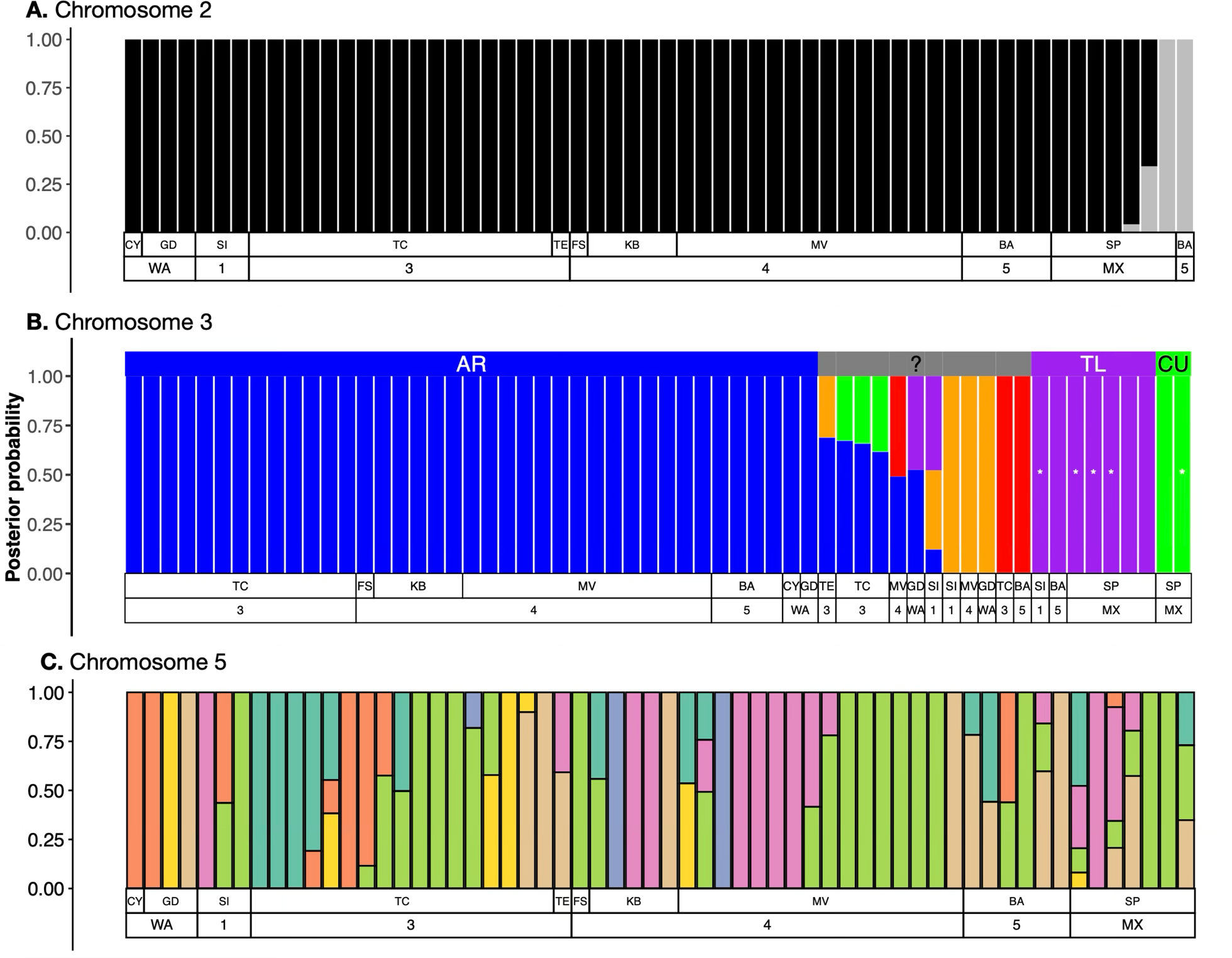
Population structure based on posterior probabilities of assignment to genotype clusters. Each bar represents one of the 60 strains, and the height of each color segment within a bar reflects the probability of assignment to a particular genetic cluster. The two rows below each graph indicate the population from which each strain was sampled and the associated ecological niche (see Figure 1). **A.** Bars show the probabilistic assignment of individuals based on chromosome 2 genetic variation to *K*=2 populations (black and gray). **B.** Probabilistic assignment of individuals to *K*=5 populations is based on chromosome 3 genetic variation, where populations likely correspond to third chromosome inversion arrangements. Blue bars correspond to the AR-assigned strains, purple bars correspond to the TL-assigned strains, and green bars correspond to the CU-assigned strains. The red and orange colors indicate additional clusters of uncertain or mixed ancestry. Asterisks show individuals that were karyotyped for specific chromosomal arrangements (Fuller et al. 2016). **C.** Bars show the probabilistic assignment of individuals based on chromosome 5 genetic variation to *K*=5 populations, each colored differently.

We hypothesized that the population structure on chromosome 3 could be explained by the third chromosome inversion polymorphism (Figure 2B). The frequencies of the third chromosome inversion arrangement differ across the six ecological niches from which we have sampled our strains (Schaeffer 2008) (Figure 1). To evaluate if the inversion polymorphism explains the chromosome 3 population structure, we attempted to assign each of the five populations to third chromosome inversion arrangements. The most common population (blue in Figure 2B) likely corresponds to the AR arrangement because AR is at high frequencies in niches 3, 4, and 5, where those strains were sampled (Figure 1). Individuals with >99% posterior probability of ancestry in the blue population were therefore assigned an AR genotype. The second most common population (purple in Figure 2B) likely corresponds to the TL arrangement because four of those strains have been karyotyped as TL homozygotes (Fuller et al. 2016). Additional individuals were assigned the TL genotype if they had >99% posterior probability assignment to the purple population. Two samples likely corresponded to the CU arrangement, one of which was previously karyotyped as homozygous for the CU arrangement (Fuller et al. 2016). These strains had >99% posterior probability of assignment to the green population (Figure 2B). The remaining 12 samples could not be unambiguously assigned to one of the arrangement genotypes because we lacked a karyotyped individual or they had mixed population assignments. Some strains with mixed assignment may be segregating for multiple chromosomal arrangements (despite being inbred). Alternatively, genetic exchange between inversion arrangements may have created haplotypes with mixed arrangement assignments (Schaeffer et al. 2024). Additional work is required to test these hypotheses for strains without an arrangement assignment.

PCA revealed a similar pattern of population structure on chromosome 3 as we found with fastSTRUCTURE (Supplementary Figure S3). PC1 explained 18.97% of the genetic variation, and strains assigned to the AR arrangement in our fastSTRUCTURE analysis were clustered separately from the remaining strains along the PC1 axis. Strains assigned to the TL and CU arrangements were also clustered together along most PC axes, but the two arrangements were separated from each other by PC8. Principal components 2–7 individually explained 7.8–10.7% of the total variance, but they did not show clear clustering patterns by any of our arrangement assignments (Supplementary Figure S3).

For chromosomes X, 4, and 5, the best fitting *K* values differed between the two criteria we used to evaluate model fit when we used a simple prior. We therefore re-ran fastSTRUCTURE with a logistic prior for these chromosomes. The optimal *K* for chromosome 5 remained >4 when we used a logistic prior, demonstrating strong evidence for population structure that may be complex because we failed to converge on a single best fitting *K* (Supplementary Table S3). We could not detect any association between the geographic origins of the strains and population assignments for chromosome 5 (Figure 2C). However, PC1 for chromosome 5 explained 20.1% of variation, and it appeared to separate the samples from Niche 3 (collected in Arizona) and most samples from other populations (Supplementary Figure S2). Below, we further explore the genetic variation on chromosome 5.

Chromosomes X and 4 had the least evidence for population structure. Using a logistic prior, at least one of our model selection criteria identified *K*=1 as the best fitting value for the X and fourth chromosomes (Supplementary Table S3). The lack of a consistent signal of *K* > 1 suggests no population structure on chromosomes X and 4. However, PCA of chromosomes 4 and X identified outlier samples that resembled those we detected for chromosome 2. In particular, PC1 explained 13.08% and 13.34% of the variation on chromosomes 4 and X, respectively, and one strain each from the BA and SP populations clustered separately from other strains (Supplementary Figure S2). These patterns provide additional evidence for a small amount of population structure along the north-south axis of the species range.

### Chromosome-specific patterns of genetic diversity

We next tested for departures from neutral expectations of heterozygosity on each chromosome using Tajima’s *D* (Tajima 1989). The lack of strong evidence for population structure across most of the genome allowed us to treat our samples as if they came from a single population in these analyses. We calculated Tajima’s *D* within 100 kb sliding windows across each chromosome for three classes of sites—intergenic, intronic, or synonymous. For the third chromosome, we also calculated *D* within individual inversion arrangements.

Consistent with previous observations (e.g., Fuller et al. 2017), we observed that *D* < 0 across *D. pseudoobscura* chromosomes (Figure 3A; Supplementary Figures S4 and S5). Regardless of which sites we analyzed, the average *D* for chromosomes X, 2, 3, and 5 fell between-0.9 and-1.0. However, chromosome 5 had a substantially higher *D* than the other chromosomes (Figure 3A). We also observed distinct troughs (*D* ≪-2) and peaks (*D* ≫ 0) in Tajima’s *D* across all chromosomes, indicating local reductions or elevations in heterozygosity. The positions of these features varied among intergenic, intronic, and synonymous sites (Figure 3A; Supplementary Figures S4 and S5), with no troughs overlapping all three categories.

**Figure 3.**
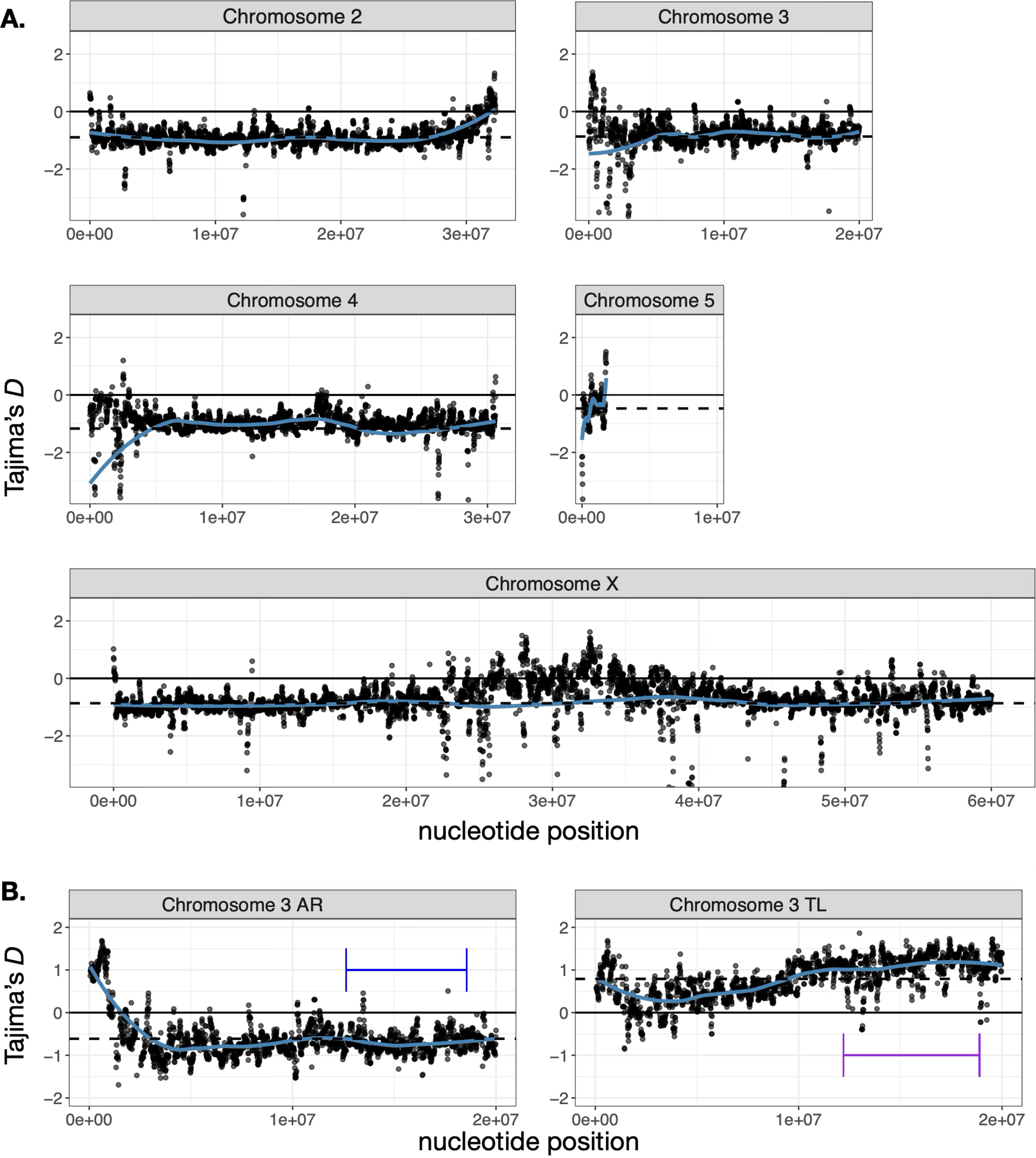
Tajima’s *D* across windows on each *Drosophila pseudoobscura* chromosome. Black dots show Tajima’s *D* values for 100 kb sliding windows (with a 10 kb step) using intergenic sites across each of the five chromosomes arms. The solid blue lines show the generalized additive models with integrated smoothness estimation (GAMs) for each chromosome. The dashed black lines show the mean Tajima’s *D* values for each chromosome. **A.** Tajima’s *D* was calculated using data from each chromosome based on all 60 strains in the dataset. **B.** Tajima’s *D* was calculated using third chromosome genetic variation from AR-assigned or TL-assigned strains. Blue and purple horizontal lines indicate the positions of the AR and TL inversions, respectively.

To further investigate regions potentially shaped by positive selection, we focused on troughs, defined as genomic windows with Tajima’s *D* <-2. This threshold is commonly used to detect regions with significantly reduced diversity, often interpreted as evidence of recent selective sweeps (Tajima 1989; Nielsen et al. 2005). Given the 100 kb window size that we used, these troughs often spanned broad genomic regions, meaning that genes within or overlapping these windows were likely affected by the same evolutionary pressures influencing local genetic variation. We specifically searched for genes located within overlapping troughs identified in at least two functional site classes. We found such overlapping troughs between intergenic and intronic sites in six windows on chromosome 3 and six windows on chromosome 4 (Supplementary Table S4). However, we did not identify any functional annotations that were significantly enriched or depleted among the genes within these regions. In addition, none of the genes located in these overlapping troughs matched previously identified candidate targets of selection in the *D. pseudoobscura* genome (Fuller et al. 2017).

We next examined the two measures of genetic diversity that are used to calculate Tajima’s *D*, average pair-wise diversity between haplotypes (*π*) and number of segregating sites (*Θ*_W_). Estimates of *π* and *Θ*_W_ varied across the lengths of chromosomes, largely in similar patterns regardless of which class of sites we used (Figure 4; Supplementary Figures S6–S10). We observed reductions in both *π* and *Θ*_W_ at one or both ends of chromosomes 2 and 4, as well as near the center of the X chromosome, likely reflecting the effects of suppressed recombination near centromeres or telomeres (Langley et al. 2012; Mackay et al. 2012). The most striking pattern emerged on chromosome 3, where genetic variation was dramatically lower on the left end of the chromosome and rose substantially higher in the right-most two thirds. The pattern on chromosome 3 is likely attributable to sampling different inversion arrangements (see below). However, because *π* and *Θ*_W_ were similarly affected across the entire genome, Tajima’s *D* did not vary as dramatically as its component genetic diversity metrics (Figure 3A).

**Figure 4.**
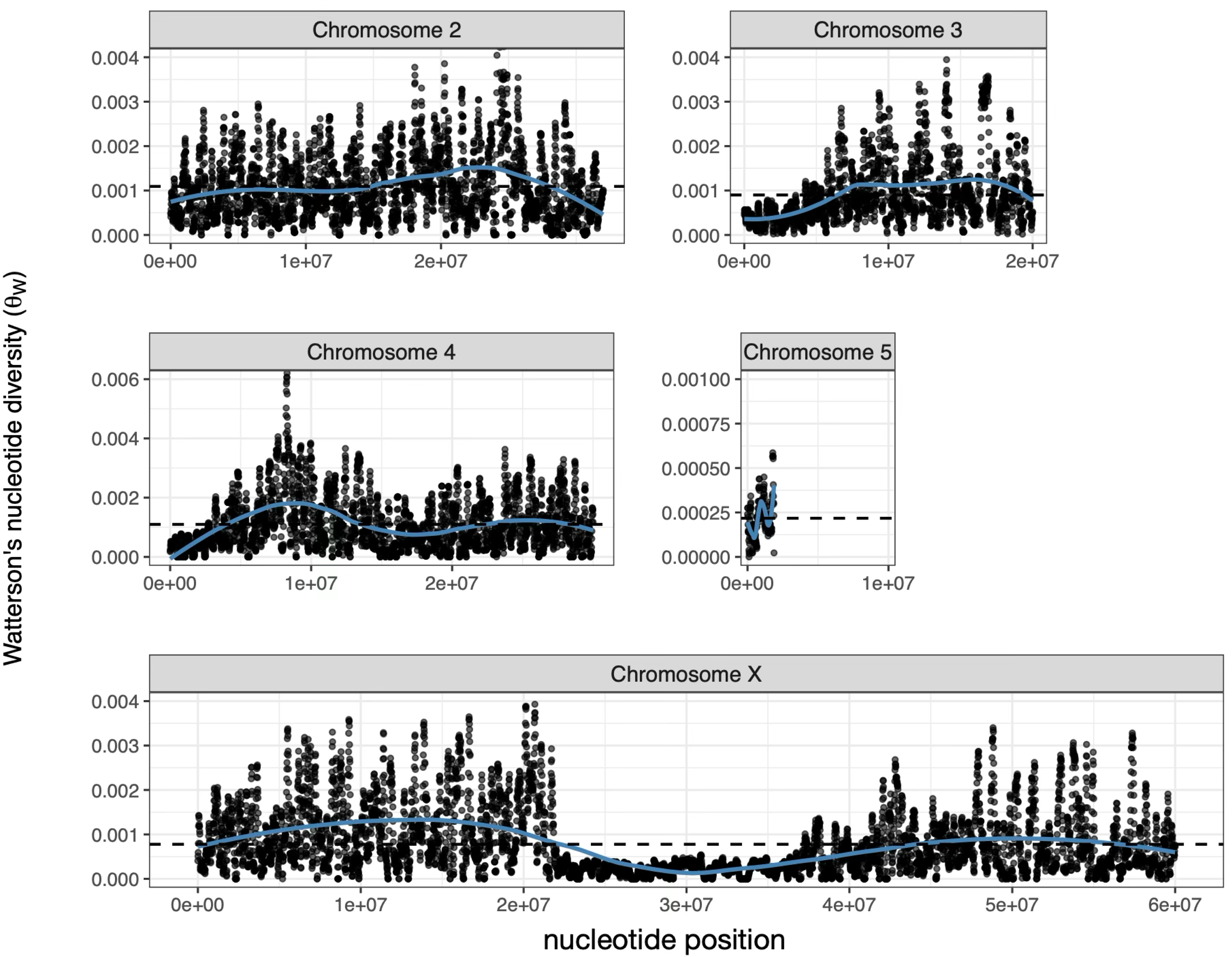
Segregating genetic variation (Watterson’s *Θ*_W_) across windows on each *Drosophila pseudoobscura* chromosome. Black dots show *Θ*_W_ for 100 kb sliding windows (with a 10 kb step) using intergenic sites across each of the five chromosomes arms. The solid blue lines show the generalized additive models with integrated smoothness estimation (GAMs) for each chromosome. The dashed black lines show the mean *Θ*_W_ for each chromosome.

### Genetic variation is affected by chromosome 3 inversions

We further investigated how the inversions affect genetic variation on chromosome 3. Most of the chromosome 3 inversions are located on the right side of the chromosome (Fuller et al. 2017), including the inversions that gave rise to the AR and TL arrangements that we hypothesize make up the majority of our samples (Figure 2). As described above, we detected elevated genetic variation on the right side of chromosome 3 (Figure 4; Supplementary Figures S6-S10). Elevated *π* is consistent with differentiation between arrangement haplotypes within the inverted regions (Navarro et al. 2000; Machado et al. 2007; Noor et al. 2007), and elevated *Θ*_W_ is consistent with sampling arrangement-specific alleles from multiple arrangements (Fuller et al. 2017). We therefore hypothesized that the inversions affected genetic variation on chromosome 3 within strains and differences between arrangement haplotypes.

To determine how the chromosome 3 inversions affected genetic variation, we separately analyzed data from strains assigned to the AR arrangement and strains assigned to the TL arrangement, based on the fastSTRUCTURE analysis (Figure 2B). Consistent with the pattern in our complete dataset, *D* < 0 along the entire length of chromosome 3 for AR-assigned strains (Figure 3B; Supplementary Figure S11). In contrast, *D* > 0 for TL-assigned strains across the entire third chromosome. We converted the chromosome 3 coordinates into the order found in the TL arrangement, and we observed the highest *D* values for TL individuals on the right half of the third chromosome, regardless of whether we used intergenic, intronic, or synonymous sites to calculate *D* (Figure 3B; Supplementary Figure S11). The elevated *D* coincides with the location of the inversion that gave rise to the TL arrangement (Figure 3B).

We then used F_ST_ for intergenic sequences as a measure of differentiation between chromosomal arrangements. Comparing AR-assigned strains against TL-assigned strains, we observed dramatically higher F_ST_ within and adjacent to the location of the inversion that gave rise to the AR arrangement (Figure 5A). We detected a less pronounced elevation with the same region of chromosome 3 when we compared AR-assigned individuals against individuals not assigned to either AR or TL (Figure 5B). Similar patterns were observed when we compared TL against AR or all other individuals using the TL coordinates for chromosome 3 (Figure 5C-D). These results provide a consistent signal that the chromosomal inversions affect genetic differentiation between third chromosome arrangements.

**Figure 5.**
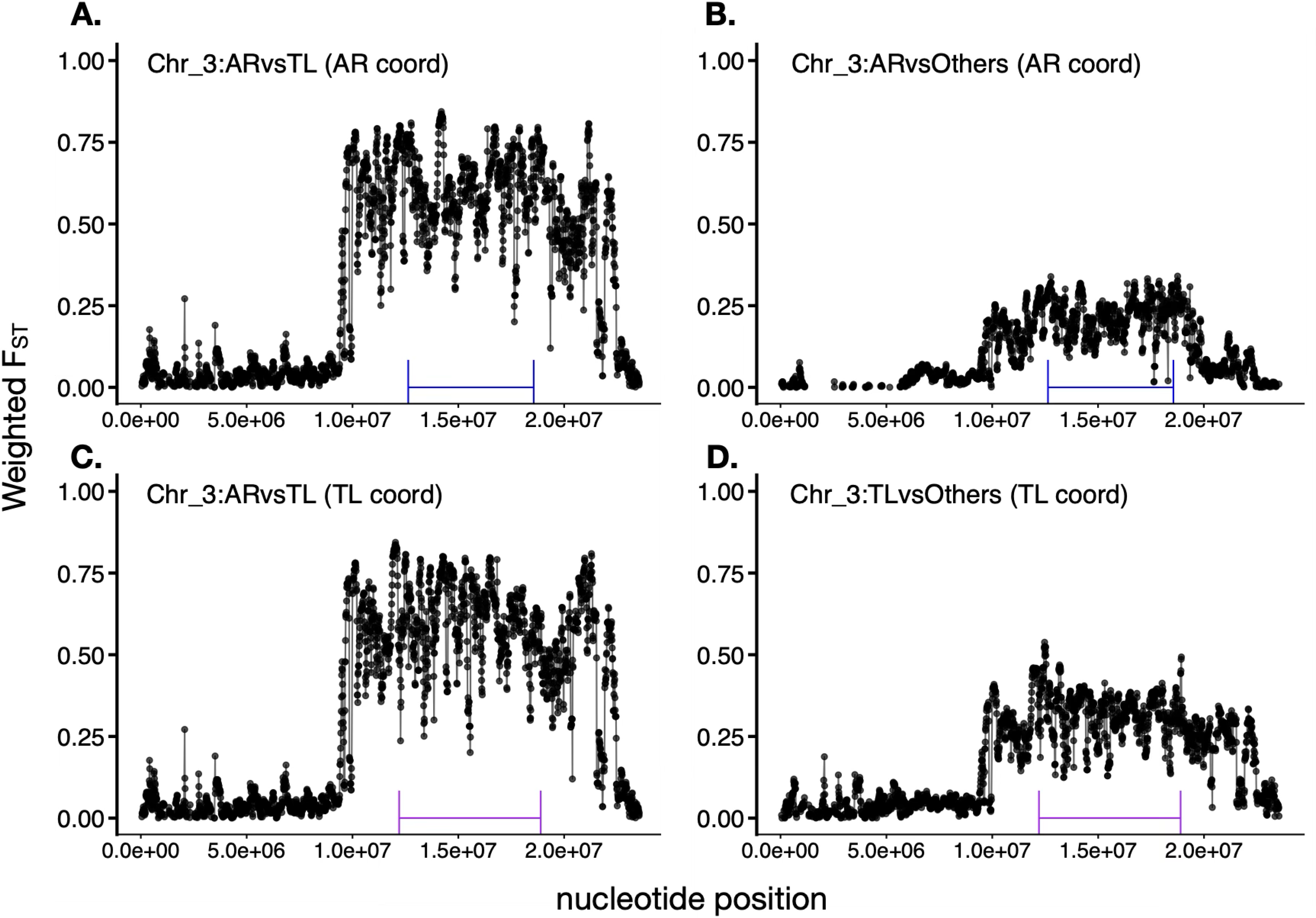
Arrangement-specific patterns of genetic differentiation across chromosome 3. Black dots show F_ST_ for 100 kb sliding windows (with a 10 kb step) using intergenic sites across chromosome 3. **A.** F_ST_ was calculated between AR-assigned and TL-assigned strains, and the AR coordinates were used for the nucleotide positions. **B.** F_ST_ was calculated between AR-assigned strains and strains not assigned to either AR or TL, with the AR coordinates for nucleotide positions. **C.** F_ST_ was calculated between AR-assigned and TL-assigned strains, and the TL coordinates were used for the nucleotide positions. **D.** F_ST_ was calculated between TL-assigned strains and strains not assigned to either AR or TL, with the AR coordinates for nucleotide positions.

## Discussion

We generated a population genomics dataset consisting of 60 *D. pseudoobscura* strains that were independently derived from flies sampled from natural populations across the species range (Figure 1). We found very little evidence for population structure across most of the genome (Figure 2A; Supplementary Table S3), but Tajima’s *D* < 0 suggests that the species likely experienced a recent expansion in size (Figure 3A). Our inferences about population structure and growth from our genome-wide data are consistent with previous findings based on small numbers of loci (Schaeffer and Miller 1992; Hamblin and Aquadro 1999; Kovacevic and Schaeffer 2000; Schaeffer et al. 2001; Machado et al. 2002; Schaeffer et al. 2003). In contrast to most of the genome, we identified strong evidence for population structure on chromosome 3 (Figure 2B), which is likely attributable to the well documented inversion polymorphism (Anderson et al. 1991; Fuller et al. 2019). We furthermore found that patterns of genetic variation differed between third chromosome inversion arrangements (Figures 3B and 4), while divergence between haplotypes appeared to be affected by the inversions (Figure 5).

We observed very little evidence for population structure across most of the genome. For example, on chromosomes 4 and X, we could not reject a model with *K*=1 (Supplementary Table S3). These findings are consistent with previous analyses of individual loci that found no evidence for population structure in *D. pseudoobscura* (Schaeffer and Miller 1992; Kovacevic and Schaeffer 2000). In contrast, we found evidence that *K*=2 for chromosome 2 (Figure 2A), suggesting some population structure that may be localized to portions of the genome. We detected a similar moderate population structure using PCA for chromosomes 2, 4, and X (Supplementary Figure S2). The secondary population we identified in these analyses is associated with a small number of strains sampled from the southernmost collection sites in our data set. These analyses are suggestive that there may be slight restrictions to north-south gene flow across the species range.

We detected substantially more evidence for population structure on chromosome 5 than most of the other chromosomes (Figure 2C). *D. pseudoobscura* chromosome 5 differs from the rest of the genome in two important ways: first, it consists of element F (i.e., the “dot” chromosome), and, second, it contains many genes that are found on the ancestral *Drosophila* Y chromosome (Larracuente et al. 2010). *Drosophila* element F is a small chromosome with fewer than 100 genes, and it has a heterochromatin environment that differs from most of the rest of the genome (Riddle and Elgin 2018). Element F is also the ancestral X chromosome of flies, and it evolved into an autosome in the lineage leading to the *Drosophila* genus (Vicoso and Bachtrog 2013). Previous work found that Tajima’s *D <* 0 across much of *D. pseudoobscura* chromosome 5 (Larracuente and Clark 2014), consistent with our results (Figure 3A). Chromosome 5 was also previously shown to have lower pairwise nucleotide diversity than much of the rest of the genome (Chang and Larracuente 2017), but we observed that both *π* and *Θ*_W_ were lower on chromosome 5 than all other chromosomes (Figure 4; Supplementary Figures S6–10). Because *Θ*_W_ is lower on chromosome 5, it has a less negative Tajima’s D when compared to other chromosomes (Figure 2). The unique distribution of genetic variation on chromosome 5, along with an extremely low recombination rate (Wang et al. 2002; Arguello et al. 2010), may be responsible for the evidence of population structure that we discovered (Figure 2C). Further work is required to investigate the causes of the unique genetic variation on chromosome 5.

The evidence for population structure on the third chromosome (Figure 2B) can likely be explained by the inversion polymorphism that segregates in natural populations of *D. pseudoobscura* (Dobzhansky and Sturtevant 1938; Anderson et al. 1975; Anderson et al. 1991). Most of the strains we sampled probably carry the AR arrangement because this is the most common third chromosome type found in the populations from which these strains were derived (Dobzhansky 1944). We could assign some of the remaining strains to one of two additional third chromosome arrangements (TL and CU) based on previous karyotyping of flies from these same isofemale lines (Fuller et al. 2016). We could not unambiguously assign the remaining strains to arrangements, but we hypothesize that some carry the PP (likely red) and CH (likely orange) arrangements based on their abundance in the populations from which they were sampled (Figures 1 and 2B). Our results provide, to the best of our knowledge, the first example of using STRUCTURE or fastSTRUCTURE to identify inversion polymorphisms in natural populations.

The lack of pronounced population structure allowed us to treat most of the genome as if it came from a single population for the purpose of testing for deviations from equilibrium. Population structure affects neutral variation in ways that can resemble some forms of selection and dampen other signals of selection (Charlesworth et al. 1997; Nordborg and Innan 2003; Wright et al. 2003; Moeller et al. 2007). Because there was such weak evidence for population structure, we examined alternative explanations for the non-neutral patterns of genetic variation that we observed.

Without strong evidence for structure on chromosomes X, 2, and 4, we are left with three interpretations of genome-wide Tajima’s *D* < 0 in *D. pseudoobscura* (Figure 3A). The first two interpretations involve the effects of natural selection, specifically background selection or selective sweeps. If background selection against deleterious mutations and/or the adaptive fixation of beneficial alleles are sufficiently common, the majority of the genome may be genetically linked to sites under selection causing a genome-wide departure from neutral expectations (Sella et al. 2009). There is indeed evidence that background selection is pervasive in the *Drosophila melanogaster* genome (Comeron 2014). However, the effects of background selection are expected to differ depending on the extent of selective constraint on linked sites (Schrider 2020). We observed similar patterns of *D* < 0 in at intergenic, intronic, and synonymous sites (Figure 3A; Supplementary Figures S4 and S5), suggesting that background selection is not responsible for the departure from neutrality. In addition, prior studies of patterns of diversity in the *D. pseudoobscura* genome demonstrated that levels of genetic variation are not correlated with recombination rates in ways that would be consistent with pervasive selection driving departures from neutrality (Kulathinal et al. 2008). We therefore conclude that *D* < 0 genome-wide in *D. pseudoobscura* is unlikely to be caused by selective sweeps or background selection.

The third interpretation of *D* < 0 is that there has been a recent increase in the effective population size of *D. pseudoobscura*. We observed that Tajima’s *D* < 0 across the genome (Figure 3A), which is consistent with previous studies of individual *D. pseudoobscura* loci or chromosomes (Hamblin and Aquadro 1999; Kovacevic and Schaeffer 2000; Schaeffer et al. 2001; Machado et al. 2002; Schaeffer 2002; Schaeffer et al. 2003; Larracuente et al. 2010; Fuller et al. 2017). When *D* < 0 across the entire genome, it suggests that demographic history (e.g., population expansion) is responsible for the pattern (Hartl and Clark 2007).

Additional evidence that *D* < 0 is caused by an increase in population size comes from X-autosome contrasts. Both arms of the *D. pseudoobscura* X chromosome (i.e., the ancestral X and neo-X) have average Tajima’s *D* values that resemble the major autosomes (Figure 3A). If selective sweeps of new recessive mutations were responsible for *D* < 0, we would expect those effects to be greater on the X than the autosomes (Harris et al. 2024). The greater effect on X-linked genes is expected because recessive beneficial mutations are exposed to selection on the X in males, who only have a single X chromosome, while autosome recessive mutations are masked (Charlesworth et al. 1987; Meisel and Connallon 2013). Not only do we not observe X-autosome differences in Tajima’s *D*, we also do not observe differences between the neo-X and ancestral X arms (Figure 3A). The lack of X-autosome differences in Tajima’s *D* is consistent with demographic effects (population expansion) driving the genome-wide *D* < 0.

Even more evidence that demographic history, as opposed to selection, drives *D* < 0 in *D. pseudoobscura* comes from X-autosome comparisons of *Θ*_W_ and *π*. In contrast to Tajima’s *D*, both *Θ*_W_ and *π* for the X chromosome are lower than the autosomes (Figure 4; Supplementary Figures S6–S10). *Θ*_W_ and *π* each scale with the product of the effective population size (*N_e_*) and mutation rate (Watterson 1975; Tajima 1989). X-autosome differences in *Θ*_W_ and *π* could thus be caused by differences in the population-level mutation rates and/or differing *N_e_*. A higher autosomal mutation rate can arise because of female-biased transmission of the X chromosome and higher mutation rates in the male germline, but evidence for this phenomenon in *Drosophila* is mixed (Bauer and Aquadro 1997; Bachtrog 2008; Wang et al. 2023). Alternatively, *N*_e_ of the X chromosome (*N*_eX_) is expected to be 75% of *N*_e_ of the autosomes (*N*_eA_) because there are 3/4 as many X chromosomes transmitted between generations, assuming equal numbers of breeding males and females (Singh et al. 2007; Vicoso and Charlesworth 2009). Both the mutational and population size effects would give rise to lower *Θ*_W_ and *π* on the X chromosome as compared to the autosomes, but the magnitude of the effects should be larger if under the population size effect (Ellegren 2007). We observed X/autosome ratios in *Θ*_W_ and *π* on the scale of 75% (Figure 4; Supplementary Figures S6–S10), consistent with the X-autosome differences being caused by *N*_eX_ < *N*_eA_. Once again, this is a demographic explanation for patterns of genetic diversity, suggesting that demographic history best explains the genome-wide average deviation from equilibrium. This supports our hypothesis that population expansion is the best explanation for Tajima’s *D* < 0 genome-wide.

Our conclusion about the prevailing effect of population expansion on genetic variation in *D. pseudoobscura* does not exclude selection also affecting genetic variation. For example, we observed distinct troughs (*D* ≪-2) and peaks (*D* ≫ 0) in Tajima’s *D* across all chromosomes, but the positions of these features varied among intergenic, intronic, and synonymous sites (Figure 3A; Supplementary Figures S2 and S3). No troughs overlapped all three categories, suggesting that selection may affect local patterns of genetic variation in a site-class-specific manner. Troughs in *D* could indicate loci that experienced a recent selective sweep, which could be explored in future work.

Additional evidence of selection comes from comparisons of polymorphism between third chromosome haplotypes assigned to different inversion arrangements. The majority of strains in our dataset were assigned to the AR arrangement (Figure 2), and Tajima’s *D* < 0 on the third chromosome for those AR-assigned strains (Figure 3). There was also a moderate elevation of *D* near the breakpoints of the AR inversion in those AR-assigned strains (Figure 3B), consistent with previous results (Fuller et al. 2017). AR is one of the youngest (most recently derived) third chromosome arrangements, and it has recently increased in frequency across much of the species’ range (Wallace et al. 2011; Fuller et al. 2017). This increase in frequency is consistent with *D* < 0 when we sample only AR-assigned strains. In addition, when we compared AR with all other arrangements, we observed elevated F_ST_ within the inverted region (Figure 5B). Elevated F_ST_ between inversion arrangements is expected for adaptively diverging arrangements (Guerrero et al. 2012).

Third chromosomes that we assigned to the TL arrangement were notable outliers, with Tajima’s *D* > 0 (Figure 3B). *D* was particularly elevated within and surrounding the location of the inversion that gave rise to the TL arrangement, consistent with previous results (Fuller et al. 2017). TL is one of the oldest inversion arrangements (Wallace et al. 2011), and TL chromosomes are rare across most of the species’ range (Dobzhansky 1944; Anderson et al. 1991). If TL has decreased in frequency, it could harbor an excess of intermediate frequency alleles (Maruyama and Fuerst 1985), which would cause *D* > 0.

Our analyses of genome-wide SNPs in *D. pseudoobscura* both confirmed previous findings and uncovered new patterns that were not observed in prior work. For example, we confirmed inferences about the effects of demographic history and third chromosome inversion arrangements on genetic variation that were made in previous studies using fewer loci, fewer individuals, and no chromosome scale reference. However, we also showed how polymorphic chromosomal inversions can be detected from signatures of population structure. Specifically, our work demonstrated how combining tests for population structure with analyses of genetic variation and differentiation can identify inversion. We additionally discovered additional outlier patterns of genetic variation on *D. pseudoobscura* chromosome 5 that should be examined in future work. The availability of our population genomics resource will facilitate such work.

## Acknowledgements

This work was supported by start-up funds from the University of Houston awarded to RPM and National Institutes of Health grant R35-GM152232 to RPM.

## Data Availability

All sequence data have been deposited in the NCBI SRA under BioProject accession PRJNA1413398. Additional data and analysis commands are available from dx.doi.org/10.18738/T8/QLQBOD.

## Supplementary Materials for A population genomic resource for *Drosophila pseudoobscura*

**Supplementary Table S1.** List of 60 *D. pseudoobscura* strains used in this study. Strains were sampled from 10 geographically distinct populations across the species’ range in North America. The four samples that did not pass filtering (see the Methods) are highlighted in red and labeled as “NO” in the “retained?” column.

| sample number | strain | pop code | location | known arrangement | retained? |
| --- | --- | --- | --- | --- | --- |
| 1 | MV-1 1108 | MV | Mesa Verde, CO | - | YES |
| 2 | MV-5 1110 | MV | Mesa Verde, CO | - | YES |
| 3 | MV-6 1111 | MV | Mesa Verde, CO | - | YES |
| 4 | MV-7 1112 | MV | Mesa Verde, CO | - | YES |
| 5 | MV-10 1114 | MV | Mesa Verde, CO | - | YES |
| 6 | MV-11 1115 | MV | Mesa Verde, CO | - | YES |
| 7 | MV-15 1116 | MV | Mesa Verde, CO | - | YES |
| 8 | MV 18 | MV | Mesa Verde, CO | - | YES |
| 9 | MV-19 1118 | MV | Mesa Verde, CO | - | YES |
| 10 | MV-21 1119 | MV | Mesa Verde, CO |  | YES |
| 11 | MV-23 1120 | MV | Mesa Verde, CO | - | YES |
| 12 | MV-25 1121 | MV | Mesa Verde, CO | - | YES |
| 13 | MV-26 1122 | MV | Mesa Verde, CO | - | YES |
| 14 | MV-28 1123 | MV | Mesa Verde, CO | - | YES |
| 15 | MV-30 1124 | MV | Mesa Verde, CO | - | YES |
| 16 | BMC-1 | BA | Bosque de Apache NWF, NM | - | YES |
| 17 | BMC-2 | BA | Bosque de Apache NWF, NM | - | YES |
| 18 | BMC-4 | BA | Bosque de Apache NWF, NM | - | YES |
| 19 | BMC-5 | BA | Bosque de Apache NWF, NM | - | YES |
| 20 | BMC-6 | BA | Bosque de Apache NWF, NM | - | YES |
| 21 | BMC-10 | BA | Bosque de Apache NWF, NM | - | NO |
| 22 | BMC-11 | BA | Bosque de Apache NWF, NM | - | YES |
| 23 | BMC-12 | BA | Bosque de Apache NWF, NM | - | NO |
| 24 | KB-2 1098 | KB | Kaibab, AZ | - | YES |
| 25 | KB-3 1099 | KB | Kaibab, AZ | - | YES |
| 26 | KB-4 1100 | KB | Kaibab, AZ | - | YES |
| 27 | KB-5 1101 | KB | Kaibab, AZ | - | NO |
| 28 | KB-6 1102 | KB | Kaibab, AZ | - | YES |
| 29 | KB-10 1105 | KB | Kaibab, AZ | - | YES |
| 30 | SPE123-1 | SP | San Pablo Etla, MX | - | YES |
| 31 | SPE123-2 | SP | San Pablo Etla, MX | TL | YES |

| sample number | strain | pop code | location | known arrangement | retained |
| --- | --- | --- | --- | --- | --- |
| 32 | SPE123-4 | SP | San Pablo Etla, MX | CU | YES |
| 33 | SPE123-5 | SP | San Pablo Etla, MX | TL | YES |
| 34 | SPE123-6 | SP | San Pablo Etla, MX | TL | YES |
| 35 | SPE123-7 | SP | San Pablo Etla, MX | - | YES |
| 36 | SPE123-8 | SP | San Pablo Etla, MX | - | YES |
| 37 | MV2-25 | MV | Mesa Verde, CO | - | YES |
| 38 | Flagstaff | FS | Flagstaff, AZ | - | YES |
| 39 | Gold-108 | GD | Goldendale, WA | - | YES |
| 40 | Gold-14B | GD | Goldendale, WA | - | YES |
| 41 | Gold-47 | GD | Goldendale, WA | - | YES |
| 42 | Tempe-5 | TE | Tempe, AZ | - | YES |
| 43 | Cheney-66 | CY | Cheney, WA | - | YES |
| 44 | IsoTucson-6 | TC | Tucson, AZ | - | NO |
| 45 | IsoTucson-7 | TC | Tucson, AZ | - | YES |
| 46 | IsoTucson-10 | TC | Tucson, AZ | - | YES |
| 47 | IsoTucson-13 | TC | Tucson, AZ | - | YES |
| 48 | IsoTucson-14 | TC | Tucson, AZ | - | YES |
| 49 | IsoTucson-20 | TC | Tucson, AZ | - | YES |
| 50 | IsoTucson-1 | TC | Tucson, AZ | - | YES |
| 51 | IsoTucson-2 | TC | Tucson, AZ | - | YES |
| 52 | IsoTucson-4 | TC | Tucson, AZ | - | YES |
| 53 | IsoTucson-8 | TC | Tucson, AZ | - | YES |
| 54 | IsoTucson-11 | TC | Tucson, AZ | - | YES |
| 55 | IsoTucson-12 | TC | Tucson, AZ | - | YES |
| 56 | IsoTucson-15 | TC | Tucson, AZ | - | YES |
| 57 | IsoTucson-16 | TC | Tucson, AZ | - | YES |
| 58 | IsoTucson-17 | TC | Tucson, AZ | - | YES |
| 59 | IsoTucson-19 | TC | Tucson, AZ | - | YES |
| 60 | IsoTucson-21 | TC | Tucson, AZ | - | YES |
| 61 | IsoTucson-22 | TC | Tucson, AZ | - | YES |
| 62 | ISOSANCRUZ-2 | SI | Santa Cruz Island, CA | - | YES |
| 63 | ISOSANCRUZ-12 | SI | Santa Cruz Island, CA | TL | YES |
| 64 | ISOSANCRUZ-13 | SI | Santa Cruz Island, CA | - | YES |

**Supplementary Table S2.** Number of SNPs per chromosome after initial filtering and number of SNPs retained after applying filtering criteria for population structure analysis. SNPs were retained if they had a genotype call rate of at least 95% (--max-missing 0.95) and a minor allele frequency of at least 5% (--maf 0.05) across the remaining strains.

| Chromosome | Initial SNPs | Retained SNPs | Percent Retained |
| --- | --- | --- | --- |
| Chromosome 2 | 504,501 | 201,653 | 39.9% |
| Chromosome 3 | 351,536 | 157,367 | 44.8% |
| Chromosome 4 | 519,967 | 195,063 | 37.5% |
| Chromosome 5 | 5,396 | 1,840 | 34.1% |
| Chromosome X | 723,865 | 275,885 | 38.1% |
| sum | 2,105,265 | 831,808 | 39.5% |

**Supplementary Table S3.**
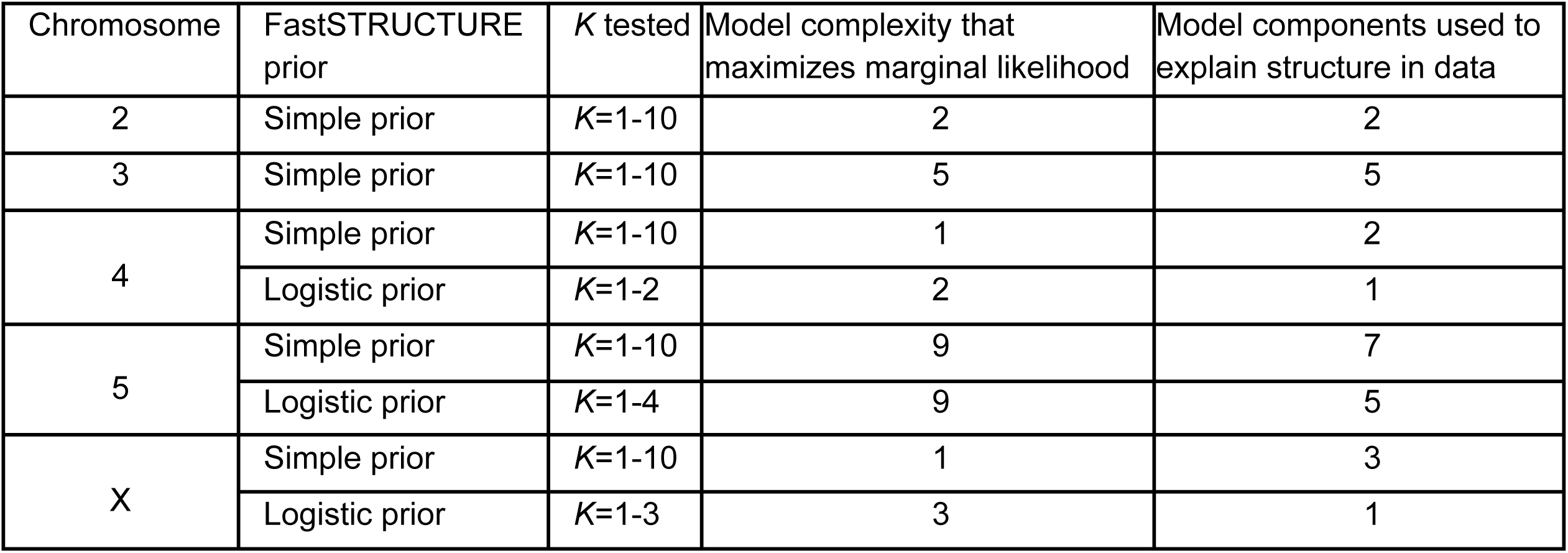
Population structure inferred by fastSTRUCTURE for each chromosome. The table shows the range of *K* values tested, along with the model complexity that maximized the marginal likelihood and the number of model components used to explain structure in the data under both the simple and logistic priors.

**Supplementary Table S4.** List of genes found in overlapping troughs of Tajima’s *D* between intergenic and intronic sites on chromosomes 3 and 4. Due to its size, Supplementary Table S3 is provided as a separate comma-separated values (CSV) file (Table_S3.csv) rather than being included in the supplemental document.

**Supplementary Figure S1.** Log10-transformed mean read depth across *Drosophila pseudoobscura* chromosomes for each strain.

**Supplementary Figure S2.** Principal component analysis showing the first two principal components (PC1 and PC2) for *Drosophila pseudoobscura* chromosomes 2, 4, 5 and X. Points are colored by population (Pop) and shaped by niche.

**Supplementary Figure S3.** Principal component analysis for *Drosophila pseudoobscura* chromosome 3. Scatterplots show pairs of principal components (PCs). Points are colored by chromosome 3 arrangement assignments (described in the main text).

**Supplementary Figure S4.** Tajima’s *D* across chromosomal windows using intronic sites in the *Drosophila pseudoobscura* genome. Black dots show Tajima’s *D* values for 100 kp sliding windows (with a 10 kp step) across each of the five chromosomes. The solid blue lines show the generalized additive models with integrated smoothness estimation (GAMs) for each chromosome. The dashed black lines show the mean Tajima’s *D* values for each chromosome.

**Supplementary Figure S5.** Tajima’s *D* across chromosomal windows using synonymous sites in the *Drosophila pseudoobscura* genome. Black dots show Tajima’s *D* values for 100 kp sliding windows (with a 10 kp step) across each of the five chromosomes. The solid blue lines show the generalized additive models with integrated smoothness estimation (GAMs) for each chromosome. The dashed black lines show the mean Tajima’s *D* values for each chromosome.

**Supplementary Figure S6.** Segregating genetic variation (Waterson’s *Θ*_W_) across windows on each *Drosophila pseudoobscura* chromosome. Black dots show *Θ*_W_ for 100 kb sliding windows (with a 10 kb step) using intronic sites across each of the five chromosomes arms. The solid blue lines show the generalized additive models with integrated smoothness estimation (GAMs) for each chromosome. The dashed black lines show the mean *Θ*_W_ for each chromosome.

**Supplementary Figure S7.** Segregating genetic variation (Waterson’s *Θ*_W_) across windows on each *Drosophila pseudoobscura* chromosome. Black dots show *Θ*_W_ for 100 kb sliding windows (with a 10 kb step) using synonymous sites across each of the five chromosomes arms. The solid blue lines show the generalized additive models with integrated smoothness estimation (GAMs) for each chromosome. The dashed black lines show the mean *Θ*_W_ for each chromosome.

**Supplementary Figure S8.** Average pair-wise differences (*π*) across windows on each *Drosophila pseudoobscura* chromosome. Black dots show *π* for 100 kb sliding windows (with a 10 kb step) using intergenic sites across each of the five chromosomes arms. The solid blue lines show the generalized additive models with integrated smoothness estimation (GAMs) for each chromosome. The dashed black lines show the mean *π* for each chromosome.

**Supplementary Figure S9.** Average pair-wise differences (*π*) across windows on each *Drosophila pseudoobscura* chromosome. Black dots show *π* for 100 kb sliding windows (with a 10 kb step) using intronic sites across each of the five chromosomes arms. The solid blue lines show the generalized additive models with integrated smoothness estimation (GAMs) for each chromosome. The dashed black lines show the mean *π* for each chromosome.

**Supplementary Figure S10.** Average pair-wise differences (*π*) across windows on each *Drosophila pseudoobscura* chromosome. Black dots show *π* for 100 kb sliding windows (with a 10 kb step) using synonymous sites across each of the five chromosomes arms. The solid blue lines show the generalized additive models with integrated smoothness estimation (GAMs) for each chromosome. The dashed black lines show the mean *π* for each chromosome.

**Supplementary Figure S11.** Tajima’s *D* across the *Drosophila pseudoobscura* third chromosome. Black dots show Tajima’s *D* values for 100 kb sliding windows (with a 10 kb step) using intronic sites (top) or synonymous sites (bottom). Tajima’s *D* was calculated using third chromosome genetic variation from either AR-assigned (left) or TL-assigned (right) strains. Blue and purple horizontal lines indicate the positions of the AR and TL inversions, respectively. The solid blue fit lines show the generalized additive models with integrated smoothness estimation (GAMs) for each chromosome. The dashed black lines show the mean Tajima’s *D*.

## References Cited

Anderson W, Dobzhansky T, Pavlovsky O, Powell J, Yardley D. 1975. Genetics of natural populations xlii. Three decades of genetic change in Drosophila pseudoobscura. Evolution. 29(1):24–36.

Anderson WW, Arnold J, Baldwin DG, Beckenbach AT, Brown CJ, Bryant SH, Coyne JA, Harshman LG, Heed WB, Jeffery DE. 1991. Four decades of inversion polymorphism in Drosophila pseudoobscura. Proc Natl Acad Sci U S A. 88(22):10367–10371.

Arguello JR, Zhang Y, Kado T, Fan C, Zhao R, Innan H, Wang W, Long M. 2010. Recombination yet inefficient selection along the Drosophila melanogaster subgroup’s fourth chromosome. Mol Biol Evol. 27(4):848–861.

Avila V, Marion de Procé S, Campos JL, Borthwick H, Charlesworth B, Betancourt AJ. 2014. Faster-X effects in two Drosophila lineages. Genome Biol Evol. 6(10):2968–2982.

Bachtrog D. 2008. Evidence for male-driven evolution in Drosophila. Mol Biol Evol. 25(4):617–619.

Bauer VL, Aquadro CF. 1997. Rates of DNA sequence evolution are not sex-biased in Drosophila melanogaster and D. simulans. Mol Biol Evol. 14(12):1252–1257.

Berdan EL, Barton NH, Butlin R, Charlesworth B, Faria R, Fragata I, Gilbert KJ, Jay P, Kapun M, Lotterhos KE, et al. 2023. How chromosomal inversions reorient the evolutionary process. J Evol Biol. 36(12):1761–1782.

Bracewell R, Bachtrog D. 2020. Complex evolutionary history of the Y chromosome in flies of the Drosophila obscura species group. Genome Biol Evol. 12(5):494–505.

Bracewell RR, Bentz BJ, Sullivan BT, Good JM. 2017. Rapid neo-sex chromosome evolution and incipient speciation in a major forest pest. Nat Commun. 8(1):1593.

Carvalho AB, Clark AG. 2005. Y chromosome of D. pseudoobscura is not homologous to the ancestral Drosophila Y. Science. 307(5706):108–110.

Carvalho AB, Koerich LB, Clark AG. 2009. Origin and evolution of Y chromosomes: Drosophila tales. Trends Genet. 25(6):270–277.

Chang CC, Chow CC, Tellier LC, Vattikuti S, Purcell SM, Lee JJ. 2015. Second-generation PLINK: rising to the challenge of larger and richer datasets. Gigascience. 4:7.

Chang C-H, Larracuente AM. 2017. Genomic changes following the reversal of a Y chromosome to an autosome in Drosophila pseudoobscura. Evolution. 71(5):1285–1296.

Charlesworth B, Coyne JA, Barton NH. 1987. The relative rates of evolution of sex chromosomes and autosomes. Am Nat. 130(1):113–146.

Charlesworth B, Nordborg M, Charlesworth D. 1997. The effects of local selection, balanced polymorphism and background selection on equilibrium patterns of genetic diversity in subdivided populations. Genet Res. 70(2):155–174.

Cicconardi F, Lewis JJ, Martin SH, Reed RD, Danko CG, Montgomery SH. 2021. Chromosome Fusion Affects Genetic Diversity and Evolutionary Turnover of Functional Loci but Consistently Depends on Chromosome Size. Mol Biol Evol. 38(10):4449–4462.

Cingolani P, Patel VM, Coon M, Nguyen T, Land SJ, Ruden DM, Lu X. 2012. Using Drosophila melanogaster as a Model for Genotoxic Chemical Mutational Studies with a New Program, SnpSift. Front Genet. 3:35.

Cingolani P, Platts A, Wang LL, Coon M, Nguyen T, Wang L, Land SJ, Lu X, Ruden DM. 2012. A program for annotating and predicting the effects of single nucleotide polymorphisms, SnpEff: SNPs in the genome of Drosophila melanogaster strain w1118; iso-2; iso-3. Fly. 6(2):80–92.

Comeron JM. 2014. Background selection as baseline for nucleotide variation across the Drosophila genome. PLoS Genet. 10(6):e1004434.

Cook DE, Andersen EC. 2017. VCF-kit: assorted utilities for the variant call format. Bioinformatics. 33(10):1581–1582.

Danecek P, Auton A, Abecasis G, Albers CA, Banks E, DePristo MA, Handsaker RE, Lunter G, Marth GT, Sherry ST, et al. 2011. The variant call format and VCFtools. Bioinformatics. 27(15):2156–2158.

Danecek P, Bonfield JK, Liddle J, Marshall J, Ohan V, Pollard MO, Whitwham A, Keane T, McCarthy SA, Davies RM, et al. 2021. Twelve years of SAMtools and BCFtools. Gigascience. 10(2):giab008.

Dobzhansky T. 1944. Chromosomal races in Drosophila pseudoobscura and Drosophila persimilis. Carnegie Inst Wash. Publ.

Dobzhansky T. 1948. Chromosomal variation in populations of Drosophila pseudoobscura which inhabit northern Mexico. Am Nat. 82(803):97–106.

Dobzhansky T, Sturtevant AH. 1938. Inversions in the chromosomes of Drosophila pseudoobscura. Genetics. 23(1):28–64.

Ellegren H. 2007. Characteristics, causes and evolutionary consequences of male-biased mutation. Proc Biol Sci. 274(1606):1–10.

Ellegren H, Galtier N. 2016. Determinants of genetic diversity. Nat Rev Genet. 17(7):422–433.

Faria R, Navarro A. 2010. Chromosomal speciation revisited: rearranging theory with pieces of evidence. Trends Ecol Evol. 25(11):660–669.

Fuller ZL, Haynes GD, Richards S, Schaeffer SW. 2016. Genomics of Natural Populations: How Differentially Expressed Genes Shape the Evolution of Chromosomal Inversions in Drosophila pseudoobscura. Genetics. 204(1):287–301.

Fuller ZL, Haynes GD, Richards S, Schaeffer SW. 2017. Genomics of natural populations: Evolutionary forces that establish and maintain gene arrangements in Drosophila pseudoobscura. Mol Ecol. 26(23):6539–6562.

Fuller ZL, Koury SA, Leonard CJ, Young RE, Ikegami K, Westlake J, Richards S, Schaeffer SW, Phadnis N. 2020. Extensive Recombination Suppression and Epistatic Selection Causes Chromosome-Wide Differentiation of a Selfish Sex Chromosome in Drosophila pseudoobscura. Genetics. 216(1):205–226.

Fuller ZL, Koury SA, Phadnis N, Schaeffer SW. 2019. How chromosomal rearrangements shape adaptation and speciation: Case studies in Drosophila pseudoobscura and its sibling species Drosophila persimilis. Mol Ecol. 28(6):1283–1301.

Garrison E, Marth G. 2012. Haplotype-based variant detection from short-read sequencing. arXiv [q-bioGN]. http://arxiv.org/abs/1207.3907.

Guerrero RF, Kirkpatrick M. 2014. Local adaptation and the evolution of chromosome fusions. Evolution. 68(10):2747–2756.

Guerrero RF, Rousset F, Kirkpatrick M. 2012. Coalescent patterns for chromosomal inversions in divergent populations. Philos Trans R Soc Lond B Biol Sci. 367(1587):430–438.

Hamblin MT, Aquadro CF. 1999. DNA sequence variation and the recombinational landscape in Drosophila pseudoobscura: a study of the second chromosome. Genetics. 153(2):859–869.

Harris M, Kim BY, Garud N. 2024. Enrichment of hard sweeps on the X chromosome compared to autosomes in six Drosophila species. Genetics. 226(4):iyae019.

Hartl DL, Clark AG. 2007. Principles of Population Genetics. Sinauer.

Korunes KL, Machado CA, Noor MAF. 2021. Inversions shape the divergence of Drosophila pseudoobscura and Drosophila persimilis on multiple timescales. Evolution. 75(7):1820–1834.

Kovacevic M, Schaeffer SW. 2000. Molecular population genetics of X-linked genes in Drosophila pseudoobscura. Genetics. 156(1):155–172.

Kulathinal RJ, Bennett SM, Fitzpatrick CL, Noor MAF. 2008. Fine-scale mapping of recombination rate in Drosophila refines its correlation to diversity and divergence. Proc Natl Acad Sci U S A. 105(29):10051–10056.

Kulathinal RJ, Stevison LS, Noor MAF. 2009. The genomics of speciation in Drosophila: diversity, divergence, and introgression estimated using low-coverage genome sequencing. PLoS Genet. 5(7):e1000550.

Langley CH, Stevens K, Cardeno C, Lee YCG, Schrider DR, Pool JE, Langley SA, Suarez C, Corbett-Detig RB, Kolaczkowski B, et al. 2012. Genomic variation in natural populations of Drosophila melanogaster. Genetics. 192(2):533–598.

Larracuente AM, Clark AG. 2014. Recent selection on the Y-to-dot translocation in Drosophila pseudoobscura. Mol Biol Evol. 31(4):846–856.

Larracuente AM, Noor MAF, Clark AG. 2010. Translocation of Y-linked genes to the dot chromosome in Drosophila pseudoobscura. Mol Biol Evol. 27(7):1612–1620.

Liao Y, Zhang X, Chakraborty M, Emerson JJ. 2021. Topologically associating domains and their role in the evolution of genome structure and function in Drosophila. Genome Res. 31(3):397–410.

Li H. 2013. Aligning sequence reads, clone sequences and assembly contigs with BWA-MEM. arXiv [q-bioGN]. http://arxiv.org/abs/1303.3997.

Machado CA, Haselkorn TS, Noor MAF. 2007. Evaluation of the genomic extent of effects of fixed inversion differences on intraspecific variation and interspecific gene flow in Drosophila pseudoobscura and D. persimilis. Genetics. 175(3):1289–1306.

Machado CA, Kliman RM, Markert JA, Hey J. 2002. Inferring the history of speciation from multilocus DNA sequence data: the case of Drosophila pseudoobscura and close relatives. Mol Biol Evol. 19(4):472–488.

Mackay TFC, Richards S, Stone EA, Barbadilla A, Ayroles JF, Zhu D, Casillas S, Han Y, Magwire MM, Cridland JM, et al. 2012. The Drosophila melanogaster Genetic Reference Panel. Nature. 482(7384):173–178.

Manat Y, Meisel R. 2026. Drosophila pseudoobscura population genomics. doi:10.18738/T8/QLQBOD. [accessed 2026 Jun 26].

Maruyama T, Fuerst PA. 1985. Population bottlenecks and nonequilibrium models in population genetics. II. Number of alleles in a small population that was formed by a recent bottleneck. Genetics. 111(3):675–689.

McGaugh SE, Heil CSS, Manzano-Winkler B, Loewe L, Goldstein S, Himmel TL, Noor MAF. 2012. Recombination modulates how selection affects linked sites in Drosophila. PLoS Biol. 10(11):e1001422.

McKenna A, Hanna M, Banks E, Sivachenko A, Cibulskis K, Kernytsky A, Garimella K, Altshuler D, Gabriel S, Daly M, et al. 2010. The Genome Analysis Toolkit: a MapReduce framework for analyzing next-generation DNA sequencing data. Genome Res. 20(9):1297–1303.

Meisel RP, Connallon T. 2013. The faster-X effect: integrating theory and data. Trends Genet. 29(9):537–544.

Meisel RP, Hilldorfer BB, Koch JL, Lockton S, Schaeffer SW. 2010. Adaptive evolution of genes duplicated from the Drosophila pseudoobscura neo-X chromosome. Mol Biol Evol. 27(8):1963–1978.

Meisel RP, Malone JH, Clark AG. 2012. Faster-X evolution of gene expression in Drosophila. PLoS Genet. 8(10):e1003013.

Moeller DA, Tenaillon MI, Tiffin P. 2007. Population structure and its effects on patterns of nucleotide polymorphism in teosinte (Zea mays ssp. parviglumis). Genetics. 176(3):1799–1809.

Muller HJ. 1940. Bearing of the *Drosophila* work on systematics. In: Huxley J, editor. The New Systematics. Clarendon Press. p. 185–268.

Navarro A, Barbadilla A, Ruiz A. 2000. Effect of inversion polymorphism on the neutral nucleotide variability of linked chromosomal regions in Drosophila. Genetics. 155(2):685–698.

Nielsen R, Bustamante C, Clark AG, Glanowski S, Sackton TB, Hubisz MJ, Fledel-Alon A, Tanenbaum DM, Civello D, White TJ, et al. 2005. A scan for positively selected genes in the genomes of humans and chimpanzees. PLoS Biol. 3(6):e170.

Noor MAF, Garfield DA, Schaeffer SW, Machado CA. 2007. Divergence between the Drosophila pseudoobscura and D. persimilis genome sequences in relation to chromosomal inversions. Genetics. 177(3):1417–1428.

Nordborg M, Innan H. 2003. The genealogy of sequences containing multiple sites subject to strong selection in a subdivided population. Genetics. 163(3):1201–1213.

Novitski E, Peacock WJ, Engel J. 1965. CYTOLOGICAL BASIS OF “SEX RATIO” IN DROSOPHILA PSEUDOOBSCURA. Science. 148(3669):516–517.

Poplin R, Ruano-Rubio V, DePristo MA, Fennell TJ, Carneiro MO, Van der Auwera GA, Kling DE, Gauthier LD, Levy-Moonshine A, Roazen D, et al. 2018. Scaling accurate genetic variant discovery to tens of thousands of samples. bioRxiv.:201178. doi:10.1101/201178. [accessed 2023 May 31]. https://www.biorxiv.org/content/10.1101/201178v3.

Pritchard JK, Stephens M, Donnelly P. 2000. Inference of population structure using multilocus genotype data. Genetics. 155(2):945–959.

Raj A, Stephens M, Pritchard JK. 2014. fastSTRUCTURE: variational inference of population structure in large SNP data sets. Genetics. 197(2):573–589.

Riddle NC, Elgin SCR. 2018. The Drosophila Dot Chromosome: Where Genes Flourish Amidst Repeats. Genetics. 210(3):757–772.

Rieseberg LH. 2001. Chromosomal rearrangements and speciation. Trends Ecol Evol. 16(7):351–358.

Sambrook J, Russell DW. 2006. Purification of nucleic acids by extraction with phenol:chloroform. CSH Protoc. 2006(1):db.prot4455.

Samuk K, Manzano-Winkler B, Ritz KR, Noor MAF. 2020. Natural selection shapes variation in genome-wide recombination rate in Drosophila pseudoobscura. Curr Biol. 30(8):1517–1528.e6.

Schaeffer SW. 2002. Molecular population genetics of sequence length diversity in the Adh region of Drosophila pseudoobscura. Genet Res. 80(3):163–175.

Schaeffer SW. 2008. Selection in heterogeneous environments maintains the gene arrangement polymorphism of Drosophila pseudoobscura. Evolution. 62(12):3082–3099.

Schaeffer SW, Bhutkar A, McAllister BF, Matsuda M, Matzkin LM, O’Grady PM, Rohde C, Valente VLS, Aguadé M, Anderson WW, et al. 2008. Polytene chromosomal maps of 11 Drosophila species: the order of genomic scaffolds inferred from genetic and physical maps. Genetics. 179(3):1601–1655.

Schaeffer SW, Goetting-Minesky MP, Kovacevic M, Peoples JR, Graybill JL, Miller JM, Kim K, Nelson JG, Anderson WW. 2003. Evolutionary genomics of inversions in Drosophila pseudoobscura: evidence for epistasis. Proc Natl Acad Sci U S A. 100(14):8319–8324.

Schaeffer SW, Miller EL. 1992. Estimates of gene flow in Drosophila pseudoobscura determined from nucleotide sequence analysis of the alcohol dehydrogenase region. Genetics. 132(2):471–480.

Schaeffer SW, Richards S, Fuller ZL. 2024. Genomics of natural populations: gene conversion events reveal selected genes within the inversions of Drosophila pseudoobscura. G3 (Bethesda). 14(10):jkae176.

Schaeffer SW, Walthour CS, Toleno DM, Olek AT, Miller EL. 2001. Protein variation in Adh and Adh-related in Drosophila pseudoobscura. Linkage disequilibrium between single nucleotide polymorphisms and protein alleles. Genetics. 159(2):673–687.

Schrider DR. 2020. Background selection does not mimic the patterns of genetic diversity produced by selective sweeps. Genetics. 216(2):499–519.

Sella G, Petrov DA, Przeworski M, Andolfatto P. 2009. Pervasive natural selection in the Drosophila genome? PLoS Genet. 5(6):e1000495.

Singh ND, Macpherson JM, Jensen JD, Petrov DA. 2007. Similar levels of X-linked and autosomal nucleotide variation in African and non-African populations of Drosophila melanogaster. BMC Evol Biol. 7:202.

Sturgill D, Zhang Y, Parisi M, Oliver B. 2007. Demasculinization of X chromosomes in the Drosophila genus. Nature. 450(7167):238–241.

Sturtevant AH, Dobzhansky T. 1936. Geographical Distribution and Cytology of “Sex Ratio” in Drosophila Pseudoobscura and Related Species. Genetics. 21(4):473–490.

Tajima F. 1989. Statistical method for testing the neutral mutation hypothesis by DNA polymorphism. Genetics. 123(3):585–595.

Van der Auwera GA, O’Connor BD. 2020. Genomics in the Cloud: Using Docker, GATK, and WDL in Terra. O’Reilly Media, Inc.

Vicoso B, Bachtrog D. 2013. Reversal of an ancient sex chromosome to an autosome in Drosophila. Nature. 499(7458):332–335.

Vicoso B, Charlesworth B. 2009. Effective population size and the faster-X effect: an extended model. Evolution. 63(9):2413–2426.

Vicoso B, Haddrill PR, Charlesworth B. 2008. A multispecies approach for comparing sequence evolution of X-linked and autosomal sites in Drosophila. Genet Res. 90(5):421–431.

Wallace AG, Detweiler D, Schaeffer SW. 2011. Evolutionary history of the third chromosome gene arrangements of Drosophila pseudoobscura inferred from inversion breakpoints. Mol Biol Evol. 28(8):2219–2229.

Wang S, Nalley MJ, Chatla K, Aldaimalani R, MacPherson A, Wei KH-C, Corbett-Detig RB, Mai D, Bachtrog D. 2022. Neo-sex chromosome evolution shapes sex-dependent asymmetrical introgression barrier. Proc Natl Acad Sci U S A. 119(19):e2119382119.

Wang W, Thornton K, Berry A, Long M. 2002. Nucleotide variation along the Drosophila melanogaster fourth chromosome. Science. 295(5552):134–137.

Wang Y, McNeil P, Abdulazeez R, Pascual M, Johnston SE, Keightley PD, Obbard DJ. 2023. Variation in mutation, recombination, and transposition rates in Drosophila melanogaster and Drosophila simulans. Genome Res. 33(4):587–598.

Watterson GA. 1975. On the number of segregating sites in genetical models without recombination. Theor Popul Biol. 7(2):256–276.

Weir BS, Cockerham CC. 1984. Estimating F-statistics for the analysis of population structure. Evolution. 38(6):1358.

Wellenreuther M, Bernatchez L. 2018. Eco-Evolutionary Genomics of Chromosomal Inversions. Trends Ecol Evol. 33(6):427–440.

White MJD. 1969. Chromosomal rearrangements and speciation in animals. Annu Rev Genet. 3(1):75–98.

Wright D. 2023. Inferring inversion breakpoint evolution in homozygotes by examining chromatin architecture in Drosophila pseudoobscura. The Pennsylvania State University. [accessed 2026 Jun 30]. https://etda.libraries.psu.edu/files/final_submissions/28925.

Wright SI, Lauga B, Charlesworth D. 2003. Subdivision and haplotype structure in natural populations of Arabidopsis lyrata. Mol Ecol. 12(5):1247–1263.

Yoshida K, Rödelsperger C, Röseler W, Riebesell M, Sun S, Kikuchi T, Sommer RJ. 2023. Chromosome fusions repatterned recombination rate and facilitated reproductive isolation during Pristionchus nematode speciation. Nat Ecol Evol. 7(3):424–439.

Zhou Q, Bachtrog D. 2012. Sex-specific adaptation drives early sex chromosome evolution in Drosophila. Science. 337(6092):341–345.

